# Raman spectra identify vancomycin-resistant phenotypes and their transcriptomic features in *Staphylococcus aureus*

**DOI:** 10.1101/2024.05.12.593718

**Authors:** Ken-ichiro F. Kamei, Reiko Okura, Koseki J. Kobayashi-Kirschvink, Yuki Katayama, Yuichi Wakamoto

## Abstract

*Staphylococcus aureus* is a pathogenic bacterium that has caused multiple epidemics linked with the emergence of new antibiotic resistance. Vancomycin is the first-line antibiotic to treat methicillin-resistant *S. aureus* (MRSA) infections. However, several types of vancomycin-non-susceptible MRSA strains have been isolated from patients to date. Rapid assessment of their resistance levels and underlying molecular profiles is crucial for preventing their spread and counteracting resistance; however, the broad resistance spectrum and the diversity of genetic changes have impeded this practice. Here, we demonstrate that the vancomycin resistance levels of various MRSA strains can be determined using dimension-reduced Raman spectra obtained from single cells. The transcriptome profiles of the different strains can also be predicted from their dimensionally reduced Raman spectra by simple linear regression. This Raman-transcriptome correspondence allows us to map the transcriptome components onto dimension-reduced Raman space and characterize groups of genes associated with different phenotypes. Furthermore, single-cell Raman spectra predicted a cell strain with greater phenotypic heterogeneity, which was confirmed by single-cell growth analysis. Overall, our results demonstrate the efficacy of Raman spectroscopy in identifying resistant phenotypes, their associated gene expression features, and intrapopulation phenotypic heterogeneity.

## Introduction

The emergence and spread of antibiotic-resistant bacteria is a huge threat to public health worldwide. *Staphylococcus aureus* is known for its ability to acquire antibiotic resistance and has caused several epidemic waves since the introduction of antibiotics (*1, 2*). Vancomycin was introduced in the 1980s to treat severe methicillin-resistant *S. aureus* (MRSA) infections (*3*). However, various vancomycin-nonsusceptible MRSA strains have been identified clinically (*3–9*), which are often associated with treatment failures (*10–13*).

Vancomycin-nonsusceptible *S. aureus* strains fall into three categories: vancomycin-resistant *S. aureus* (VRSA), vancomycin intermediately-resistant *S. aureus* (VISA), and heterogeneous vancomycin intermediately-resistant *S. aureus* (hVISA) (*3,8,9*). In particular, VISA and hVISA prevalence has been increasing recently (*8, 9*). The minimum inhibitory concentration (MIC) for VISA is 4–8 *µ*g/mL, which is higher than that of vancomycin-susceptible MRSA (VSSA) strains (≤2 *µ*g/mL) but lower than that of VRSA (≥16 *µ*g/mL). hVISA strains are often judged as VSSA by the standard MIC test, but they contain small subpopulations with the VISA-level vancomycin resistance and could lead to treatment failures (*5, 14, 15*). hVISA strains are considered to be in intermediate evolutionary steps from VSSA to VISA (*15, 16*). As VISA and hVISA prevalence raises growing concern about longer hospitalization and higher risk of persistent infection (*8, 9*), it is critical to swiftly identify the resistant phenotypes of bacteria infecting patients and treat infections with appropriate drug regimens.

The methicillin resistance of MRSA is predominantly caused by the acquisition of the *mecA* or *mecC* gene on a mobile genetic element (*2, 3, 15, 17, 18*). However, the mechanisms behind vancomycin-nonsusceptibility are not yet fully understood and are believed to be diverse (*3, 9, 15,16,19–24*). Genetic analyses have suggested that multi-step mutations at various genetic loci can increase vancomycin resistance levels (*15, 16, 22, 23*). Consequently, it remains challenging to identify the vancomycin resistance levels through targeted gene analysis.

Despite the diverse mutational profiles, it is suggested that physiological characteristic changes such as cell wall thickening and slow growth are prevalent amongst vancomycin-nonsusceptible MRSA strains (*19,20,24–26*). Therefore, analyzing their global molecular composition and gene expression profiles might allow us to identify their vancomycin-resistance levels. Matrix-assisted laser desorption ionization time-of-flight mass spectrometry (MALDI-TOF MS) combined with a machine-learning classification can distinguish different pathogenic bacteria and has also been applied to identify MRSA and different phenotypes of vancomycin-nonsusceptible MRSA strains (*27–29*). However, MALDI-TOF MS requires culturing cells to obtain microbial biomass (for bacteria, 10^4^ to 10^6^ colony forming unit) for analysis (*27*). Furthermore, identifying the comprehensive molecular signatures of targeted cells based on the MALDI-TOF MS spectra remains intractable. In addition, due to the population-based approach of the analysis, it is intractable to unravel single-cell-level phenotypic heterogeneity within a bacterial population.

Raman microscopy also provides spectral information reflecting the comprehensive molecular composition of single cells (*30–35*) because a Raman spectrum from a cell is, in principle, the linear superposition of the spectra of its constituent biomolecules. The complexity of the molecular composition of cells and severe spectral overlap make it nearly impossible to comprehensively decompose a cellular Raman spectrum and determine the abundances of different cellular components (*30–32*). However, it is important to note that most changes in gene expression and metabolic profiles are inherently global and modular (*36–43*). Therefore, without finely and precisely decomposing the spectra and assigning peaks to molecular species, the correspondence between Raman spectra and comprehensive molecular profiles may well be unraveled. Indeed, recent studies have shown that global changes in the entire fingerprint region of cellular Raman spectra correlate with global gene expression profiles for environmentally perturbed cells (*33, 44, 45*). Consequently, transcriptomic and proteomic profiles can be inferred from Raman spectra thanks to the intrinsic low-dimensional constraints on changes in global gene expression patterns (*33, 44, 45*). However, most Raman applications have focused on the correlations between the changes in specific Raman spectral peaks and the changes in the abundance of specific biomolecules in cells, or have employed machine learning approaches to distinguish microbial species or cell types without clarifying the underlying molecular profiles. As a result, the correspondence between Raman spectra and system-level molecular profiles of cells has remained largely unexplored in Raman spectroscopy applications.

This report first demonstrates that dimension-reduced Raman spectra of VSSA, hVISA and VISA strains under a common culture condition can predict vancomycin-resistance phenotypes. We then reveal that global transcriptomic profiles are also inferable from the low-dimensional Raman spectra, demonstrating that the Raman-transcriptome correspondence holds even for genetically distinct cells. Using the Raman-transcriptome correspondence, we propose a method for representing transcriptome components with Raman axes that correlate with different phenotypic features. This method identifies several gene sets associated with vancomycin resistance and virulence. Finally, we show that the variability of single-cell Raman spectra can be used to interrogate the phenotypic heterogeneity of different strains, a clinically important phenotypic feature of the hVISA strains.

## Results

### Raman spectra distinguish cellular states of MRSA strains with distinct vancomycin susceptibility

In this study, we analyzed nine MRSA strains. Of these, three strains (N315, Mu3, and Mu50) are clinical isolates with distinct susceptibility to vancomycin (*5*), while the other six (ΔIP, ΔIP1–ΔIP5) are strains created by introducing specific mutations or by selection under rifampicin exposure (*16*). Vancomycin susceptibility tests performed in (*16*) classified N315 and ΔIP as VSSA, Mu3, ΔIP1, and ΔIP2 as hVISA, and Mu50 and ΔIP3–ΔIP5 as VISA on the basis of population measurements such as MIC tests (Fig. 1A).

**Figure 1.**
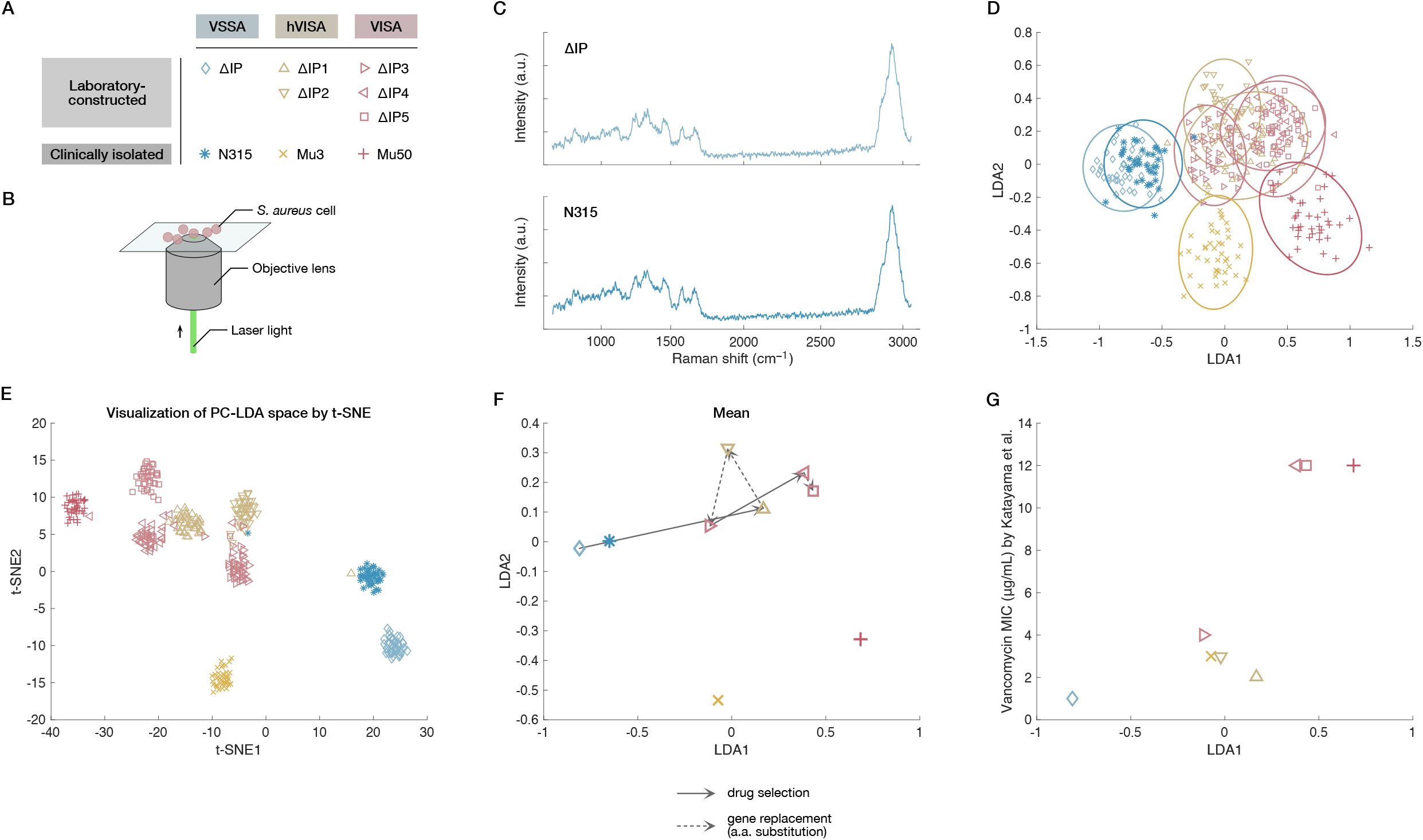
Single-cell Raman spectra of MRSA strains and their correlation with vancomycin resistance. **(A)** Nine MRSA strains analyzed in this study. Three strains are clinical isolates, while the other six are strains constructed in a laboratory (*16*). Vancomycin susceptibility of these nine strains are classified into VSSA, hVISA, and VISA. Groups to which the strains used in this study belong were judged by (*16*). Colors and markers in this figure are used in some of the other figures throughout this paper. **(B)** Schematic figure representing our Raman measurements. Raman spectra were obtained from single cells placed on a synthetic quartz slide glass by focusing laser light on each targeted cell. **(C)** Representative single-cell Raman spectra. One is from ΔIP (top), and the other is from N315 (bottom). **(D)** Cellular Raman spectra in LDA space. LDA reduces the dimensionality of the spectra to 8 (= 9 − 1). Each point represents a spectrum from a single cell. Each ellipse shows the 95% concentration ellipse. Projection to the plane spanned by the first two axes (LDA1 and LDA2) is shown. **(E)** t-SNE visualization of the eight-dimensional LDA space. **(F)** The population average of the single-cell Raman spectra of each strain in the LDA space. The arrows indicate the changes of the averages along the strain construction and selection steps in (*16*). **(G)** Correlation between the positions of cellular Raman spectra along the LDA1 axis and vancomycin resistance levels measured by (*16*), in which MIC of N315 was not reported. Pearson correlation coefficient, *r* = 0.81 *±* 0.15.

We cultured these strains in brain heart infusion (BHI) medium and sampled cells in exponential phase for Raman measurements. Raman spectra of 48 single cells per strain were acquired using a Raman microscope (Fig. 1B, 1C and S1). To eliminate systematic errors in weak spontaneous Raman signals, the spectra were acquired in three biological replicates on different dates. We focused on the fingerprint region (700 to 1,800 cm^−1^) of the Raman spectra in the analysis, where the signals from biomolecules such as proteins and metabolites are mainly observed.

We applied principal component-linear discriminant analysis (PC-LDA) to the Raman spectra to determine whether they could distinguish differences in cellular states among the strains (*33, 45, 46*). The Raman spectra were classified in a low-dimensional (eight-dimensional) space (Fig. 1D). In this LDA space, each point represents a Raman spectrum from an individual cell. The results showed that the points for the cells in each strain were closely positioned and clustered in this LDA space (Fig. 1D). Visualizing the eight-dimensional Raman spectra by embedding them in a two-dimensional space using t-distributed stochastic neighbor embedding (t-SNE) confirmed the cluster separation (Fig. 1E). Therefore, the Raman spectra can distinguish the distinct cellular states of the nine MRSA strains.

Notably, the positions along the first LDA axis (LDA1) correlated with their vancomycin resistance levels (Pearson correlation coefficient *r* = 0.81 *±* 0.15, Fig. 1F and 1G). Therefore, PC-LDA detects the Raman spectral changes correlated with the vancomycin resistance levels as the dominant differences. Additionally, the second LDA axis (LDA2) recognizes the differences between the clinically isolated vancomycin-nonsusceptible strains (Mu3 and Mu50) and the constructed vancomycin-nonsusceptible strains (ΔIP1–ΔIP5) (Fig. 1F). Therefore, the result shows that, even among the MRSA strains with equivalent vancomycin resistance levels, there exist differences in their molecular profiles detectable by Raman spectra.

### Raman spectral differences among the MRSA strains are explainable by their transcriptomes

We next investigated whether the differences in Raman spectra of nine MRSA strains can be explained by their gene expression profiles. To test this, we gathered transcriptome data of these strains cultured in the BHI medium from public databases (*16, 21, 26*) and examined their correspondences with the Raman data. We hypothesized a linear correspondence

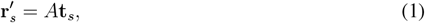

where **r***′* _*s*_ is the population average of the eight-dimensional Raman points of strain *s* (Fig. 1F), and **t**_*s*_ is the 2,628-dimensional transcriptome vector of the same strain containing the expression levels of 2,628 genes. *A* is the strain-independent matrix that maps the transcriptomes onto the Raman LDA space.

We conducted leave-one-out cross-validation (LOOCV) to verify the linear correspondence between Raman and transcriptome data (*33*). First, we estimated matrix *A* by using the Raman and transcriptome data of eight strains through partial least squares regression (PLS-R) (*47*). We then applied this estimated matrix to the transcriptome data of the excluded strain (test strain) to predict an eight-dimensional Raman point, represented as 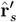, which was then compared to the actual population average point, represented as **r***′* _*s*_ (Fig. 2A). We repeated this process by excluding one strain at a time and evaluated the overall estimation error using predicted residual error sum of squares (PRESS), calculated as 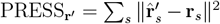. A small PRESS value validates the assumption of linear correspondence between the Raman and transcriptome data. We quantitatively evaluated the resulting PRESS_**r**_*′* value through a permutation test (*33*).

**Figure 2.**
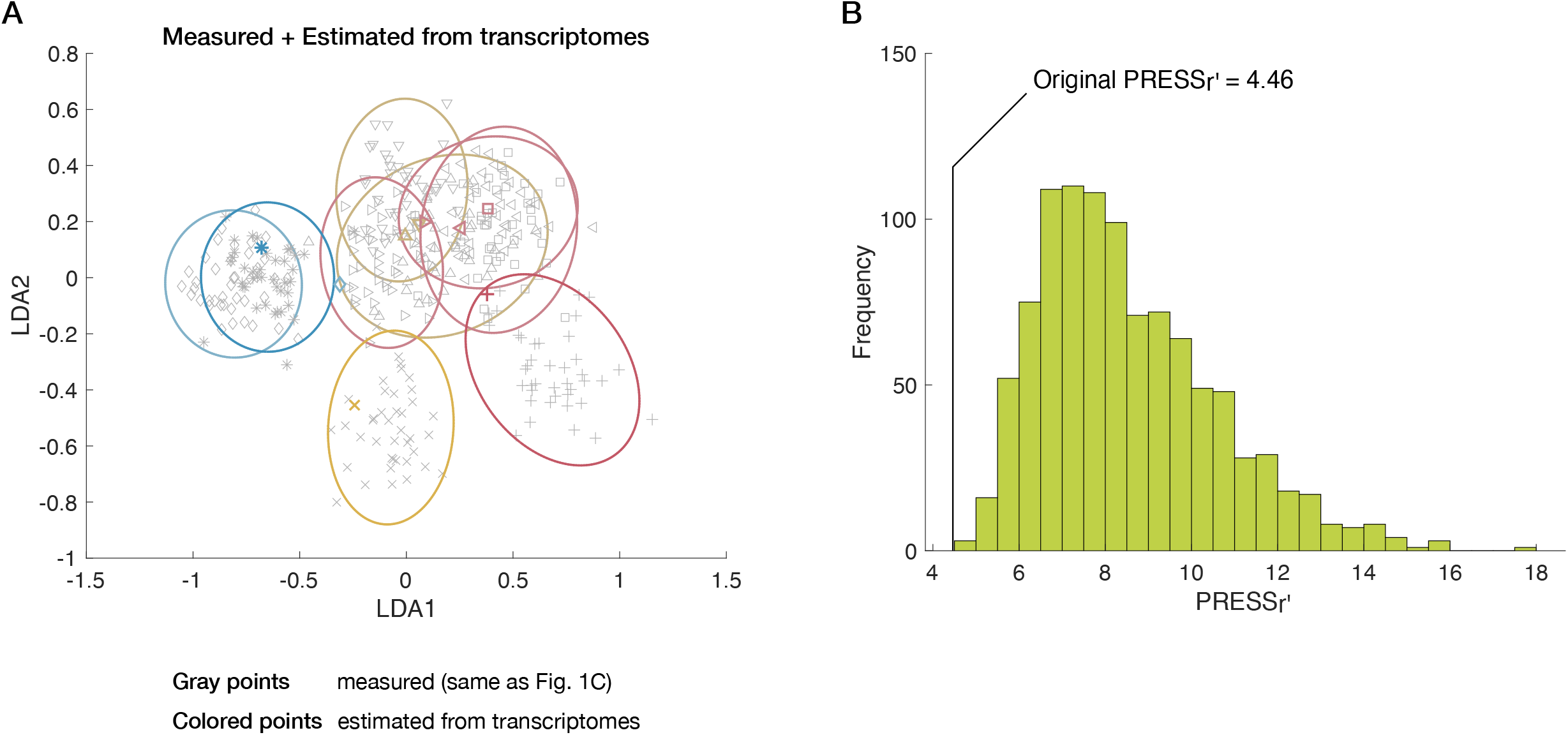
Transcriptome profiles of different MRSA strains can explain their Raman spectral differences. **(A)** Estimation of Raman spectra in the LDA space from transcriptomes. Thick colored points represent Raman spectra estimated from the transcriptomes assuming the linear correspondence (Eq. 1). **(B)** Histogram of PRESS_**r**_*′* values of 1,000 randomly permuted data. The PRESS_**r**_*′* of the original dataset is 4.46. The *p*-value was 9.99 *×* 10^−4^ (no occurrence of smaller PRESS_**r**_*′* was observed in the 1,000 permuted datasets).

We created 1,000 datasets in which we randomly permuted the correspondences between the Raman and transcriptome data, calculated the PRESS_**r**_*′* values for these false datasets, and compared them with the original PRESS_**r**_*′* value. The result confirmed a significantly small overall estimation error (PRESS_**r**_*′* = 4.46, *p* = 9.99 *×* 10^−4^, permutation test, Fig. 2B). This result therefore suggests that the observed Raman spectral differences in the LDA space can be explained by the differences in the gene expression profiles of these MRSA strains.

### Global transcriptome profiles can be inferred from low-dimensional Raman data

Next, we asked whether transcriptome profiles of different strains could be inferred from their lowdimensional Raman spectra in LDA space. To address this question, we examined another linear correspondence

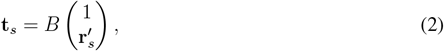

where *B* is the strain-independent coefficient matrix that converts the Raman spectral points of each strain in the LDA space into 2,628-dimensional transcriptome profile data (*45*). We again conducted LOOCV to verify this linear correspondence. We excluded the Raman and transcriptome data of one strain and estimated the coefficient matrix *B* using ordinary linear regression with the data of the other eight strains (see Materials and Methods). Then, we predicted the transcriptome profile of the excluded strain from its Raman spectrum **r***′*_*s*_ as 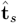 from Eq. 2.

Noticing that transcriptome profiles of different strains are also strongly correlated (Fig. S2), we calculated the PRESS for the transcriptome data as 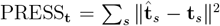 to quantify the overall estimation error. A permutation test with randomly permuted 1,000 datasets found a significantly small overall estimation error (PRESS_**t**_ = 1.09 *×* 10^4^, *p* = 9.99 *×* 10^−4^, Fig. 3C). We also quantified the overall estimation error using other distance metrics and consistently found significantly small estimation errors (Table S1). Therefore, differences in the Raman spectra of the MRSA strains contain the information that allows the prediction of global gene expression differences of these strains. Note that the transcriptome inference was performed using the low-dimensional LDA Raman data, which correlated with vancomycin MIC levels; the high precision of the Raman-transcriptome correspondence suggests a constraint on the global transcriptomic changes associated with the vancomycin resistance of the genotypically distinct MRSA strains.

**Figure 3.**
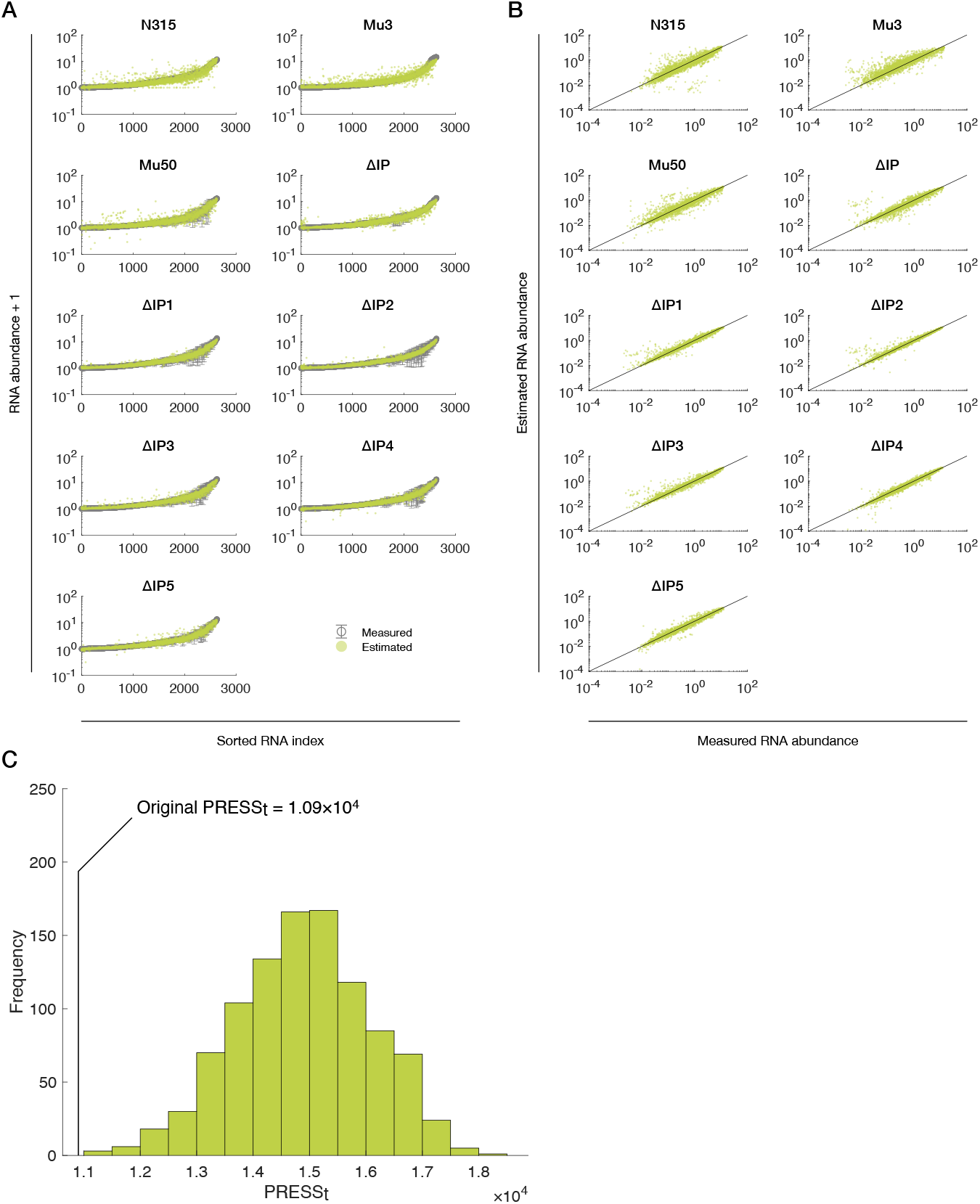
Inference of global transcriptome profiles from cellular Raman spectra. **(A and B)** Estimation of transcriptomes from Raman spectra in the LDA space. In (A), grey points represent the measured RNA abundance (average of the replicate measurements; error bar, SE), which are sorted along the horizontal axis. Green points represent the RNA abundance estimated from Raman spectra. (B) is scatterplots between the measured and the estimated RNA abundances. Straight lines represent *x* = *y*. **(C)** Histogram of PRESS_**t**_ values of 1,000 randomly permuted data. The PRESS_**t**_ of the original dataset is 1.07 *×* 10^4^. The *p*-value was 9.99 *×* 10^−4^ (there was no occurrence of smaller PRESS_**t**_ in the 1,000 permuted datasets).

### Raman-based representations of transcriptome components can identify gene groups linked to vancomycin resistance and virulence

The linear correspondence between transcriptome profiles and Raman spectra in the LDA space (Eq. 2 and Fig. 3) allows us to analyze the dependence of each gene on different LDA axes (*45*). Importantly, the major LDA axes are strongly correlated with different phenotypic features, such as vancomycin resistance levels and strain origin (clinically isolated or laboratory-constructed, Fig. 1D, 1F and 1G). Thus, the dependence of each gene on the LDA axes may help us to identify groups of genes associated with different phenotypes.

From Eq. 2, the expression level of gene *i* in strain *s* is expressed as

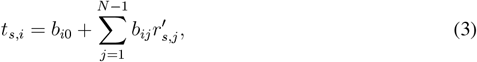

where *t*_*s,i*_ is the *i*-th element of **t**_*s*_, *b*_*i*0_ *b*_*i*1_ *·· · b*_*iN*−1_ is the *i*-th row of *B, r′* _*s, j*_ is the *j*-th element of the Raman LDA vector **r***′* _*s*_, and *N* = 9 is the number of strains in the analysis. Since *b*_*i*0_ represents the basal expression level of each gene, relative change in the expression level of gene *i* in strain *s* is

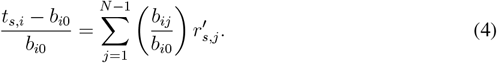

Therefore, the quantity 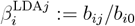 determines the dependence of gene *i* on the *j*-th LDA axis.

Figure 4 shows the plot of the first and second normalized coefficients 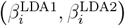 for all genes in the transcriptome data. Most genes are distributed near the center, but we can find a number of genes deviating significantly from the center. Since the LDA1 axis is linked to the vancomycin resistance, we first examined the expression patterns of the genes with high 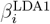 values (Table 1) and found that all of these genes are highly expressed in the VISA strains (Fig. 4). Conversely, genes with low 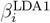 values are highly expressed in the vancomycin-susceptible strains but suppressed in the VISA strains (Fig. 4). Among the genes with high 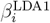 values, those with low 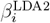 values are exclusively expressed in the clinically isolated strain Mu50, while those with high 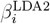 values are exclusively expressed in the laboratory-constructed strains ΔIP4 and ΔIP5 (Fig. 4). Genes with the 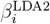 values close to zero are commonly expressed in both clinically isolated and laboratory-constructed strains (Fig. 4). These expression patterns correspond to the Raman spectral patterns in the LDA space (Fig. 1D and 1F). Thus, representing the transcriptome components using the low-dimensional Raman LDA axes allows us to distinguish genes linked with different phenotypes by corresponding Raman positions.

**Figure 4.**
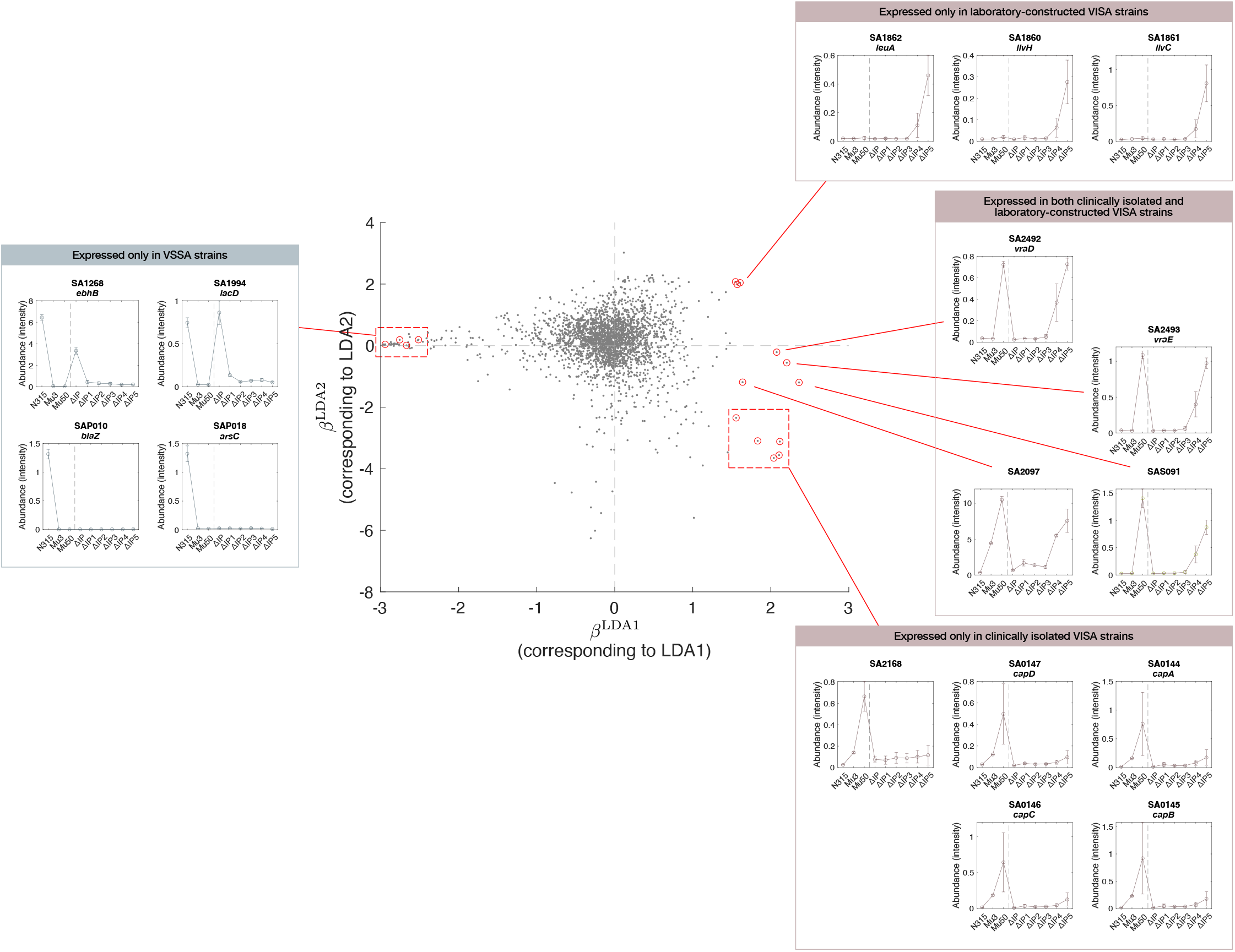
Mapping transcriptome components on dimension-reduced Raman space identifies genes associated with distinct phenotypes. All genes in the transcriptome data are represented on a plane where the coordinates of each gene *i* are its first and second normalized coefficients 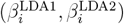. Each gray dot corresponds to a gene. All genes with 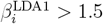 and several representative genes with 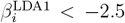 are highlighted by red circles. Expression patterns of these genes are also shown (average of the replicate measurements; error bar, SE). Since normalized coefficients 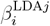 determines the dependence of gene *i* on the *j*-th LDA axis, the first and second axes (*β*^LDA1^ and *β*^LDA2^) in this figure correspond to LDA1 and LDA2 axes, respectively (Fig. 1D and 1F). For example, Raman spectra of VISA strains are placed on the right side of the LDA1-LDA2 plane in Fig. 1D and 1F. Correspondingly, genes highly expressed in the VISA strains are placed on the right side of *β*^LDA1^-*β*^LDA2^ plane in this figure.

Some of the genes identified as having high 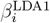 values were previously suggested to contribute to increased vancomycin resistance and virulence in *S. aureus*. Four genes—*vraD, vraE*, SAS091, and SA2097—were listed as highly expressed in both clinically isolated and laboratory-constructed VISA strains (Table 1). *vraD, vraE*, and SAS091 are ABC transporter genes that are known to be upregulated in VISA strains (*48, 49*). SA2097 encodes a peptidoglycan hydrolase component (*50, 51*). *S. aureus* possesses at least 18 different peptidoglycan hydrolases that facilitate growth and division (*50*). Since vancomycin targets the final steps of peptidoglycan and cell wall biosynthesis, high expression of SA2097 may enable the *S. aureus* cell to circumvent the action of the drug and survive. While the functions of these genes have not been fully characterized, their co-expression may be considered as a strong indicator of VISA-level vancomycin resistance.

Four of the five genes that are highly expressed exclusively in the clinically isolated VISA strain Mu50 are related to capsular polysaccharide synthesis (Table 1). It has been suggested that these genes enhance virulence by enabling *S. aureus* cells to withstand phagocytosis and killing by polymorphonuclear phagocytes (*52–54*). Thus, while the upregulation of these genes may not directly contribute to vancomycin non-susceptibility, it may be indispensable for pathogenesis.

The genes that are highly expressed exclusively in laboratory-constructed VISA strains (*ilvH, leuA*, and *ilvC*) are all in the same operon and are involved in the biosynthesis of branched-chain amino acids (BCAAs) (*55*). Thus, the coordinated upregulation of these genes may alter BCAA levels in these strains. BCAAs act as signaling molecules that can affect a number of metabolic and virulence genes by binding to the global regulatory protein CodY (*56*). However, the relevance of these changes to vancomycin non-susceptibility and virulence in these VISA strains remains unclear.

### The clinically isolated hVISA Mu3 strain is physiologically heterogeneous

Previous research has demonstrated that the hVISA Mu3 strain contains subpopulations that have the same level of resistance to vancomycin as VISA (*5*). This suggests that Mu3 is physiologically more diverse among clonal cells than those of N315 and Mu50.

In line with this picture, we noticed that the Raman spectra of the Mu3 strain were more heterogeneous among individual cells than those of the clinically isolated VSSA and VISA strains. We evaluated the standard deviations of normalized cellular Raman spectra at different wavenumbers among the clinically isolated MRSA strains (N315, Mu3, and Mu50), finding that the variations were consistently larger in Mu3 than those of N315 and Mu50 (Fig. 5A). This result suggests that the Mu3 cells are more heterogeneous in their molecular composition at the single-cell level.

**Figure 5.**
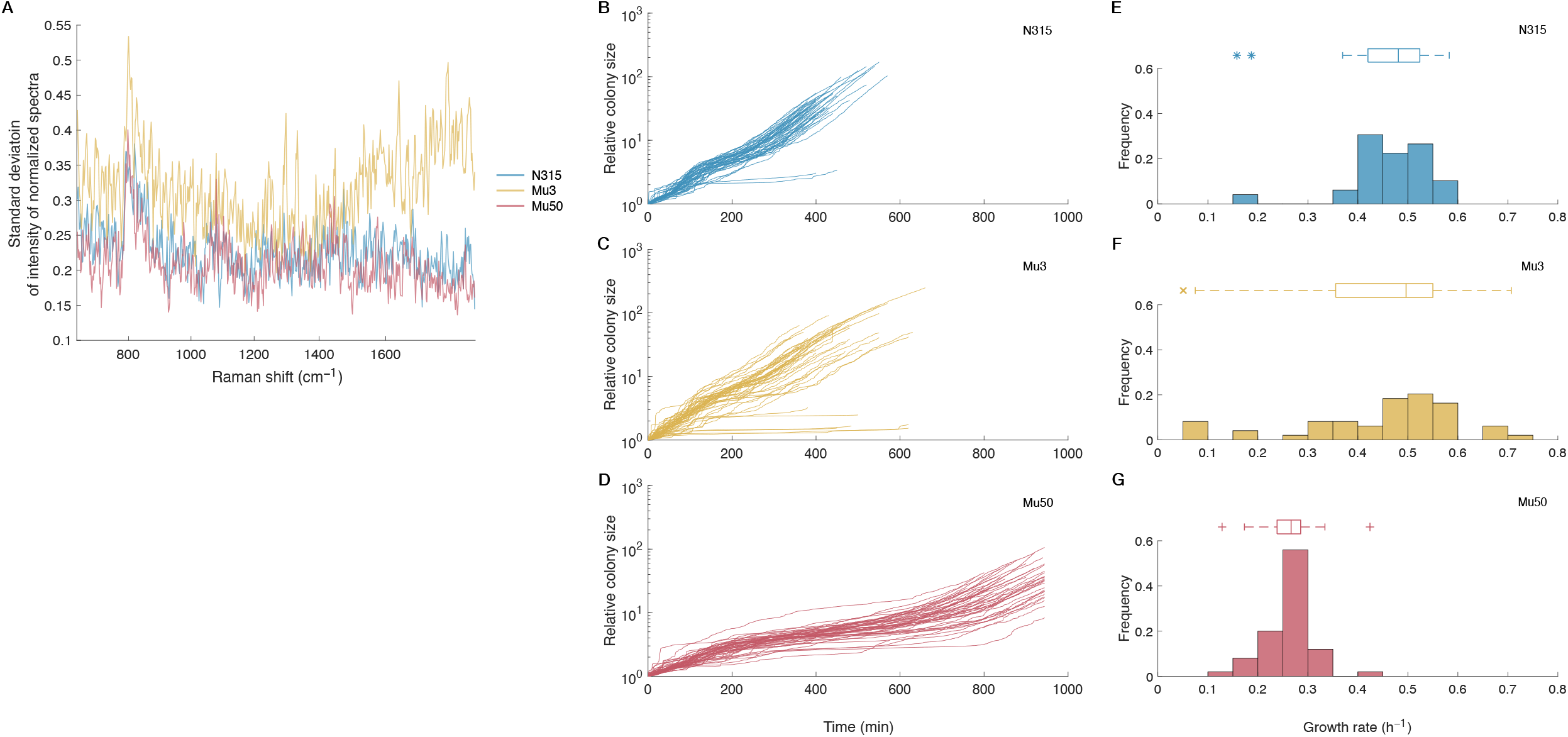
Heterogeneity of clinically isolated hVISA strain Mu3. **(A)** Comparison of standard deviations of normalized single-cell Raman spectra among the clinically isolated VSSA, hVISA, and VISA strains. **(B–D)** Microcolony growth of the three clinically isolated strains on BHI agar. Single lines represent relative size growths of different single colonies. **(E–G)** Growth rate distributions of the microcolonies on BHI agar shown in Fig. 5B–5D.

To ask whether the heterogeneity recognized by their Raman spectra is physiologically relevant, we observed the growth and microcolony formation of the three clinically isolated strains on the BHI agar by time-lapse microscopy (Movies S1–S3). As expected, the growth curves of Mu3 were more heterogeneous than those of N315 and Mu50 among the microcolonies grown from different individual cells (Fig. 5B–5D). The coefficient of variation (CV) of the growth rate of Mu3 was 36%, significantly higher than those of N315 and Mu50, which were both 18% (Fig. 5E–5G). The mean growth rate of Mu50 was smaller than that of N315 (0.47 h^−1^ for N315 and 0.26 h^−1^ for Mu50), which corroborates the slow-growing trait of the VISA strain. Although the median of the growth rate of Mu3 is close to that of N315, Mu3’s growth rate distribution is skewed with a heavy left tail, indicating the presence of subpopulations in the Mu3 strain growing as slowly as the Mu50 strain (Fig. 5E–5G).

The fact that cell-to-cell heterogeneity of Mu3 is consistently larger in both Raman spectra and growth supports that this clinically isolated hVISA strain is physiologically more diverse among genetically identical cells. Furthermore, the results suggest that phenotypic heterogeneity in cellular population can be interrogated spectroscopically by evaluating their Raman spectral heterogeneity.

## Discussion

Vancomycin resistance undermines the treatment efficacy of MRSA infection. Inappropriate administration of vancomycin could result in generating and enhancing the resistance of bacteria. Therefore, it is important to quickly detect vancomycin resistance to make the right choices in drug regimens and prevent the emergence and spread of vancomycin-nonsusceptible MRSA strains.

This study shows that the global transcriptome profiles and vancomycin resistance levels of the MRSA strains can be identified by their Raman spectral patterns (Fig. 1G, 3A, and 3B). Cellular Raman spectra can be obtained by exposing cells to light without the need for staining or tagging. Furthermore, unlike the MALDI-TOF MS-based method, it is not necessary to culture cells to target bacterial cells for Raman measurements, as long as they are recognizable in specimens. As a result, Raman microscopy has significant potential for quickly uncovering antibiotic resistance of bacteria in patients.

Previous studies have shown that cellular Raman spectra obtained under different culture conditions can be used to infer transcriptomic and proteomic differences (*33, 44, 45*). The present study shows that transcriptomic differences among the MRSA strains with different genotypes under identical culture conditions can also be inferred from their Raman spectral patterns (Fig. 3A and 3B). Therefore, Raman-based inference of gene expression profiles applies not only to external environmental differences but also to internal genotypic differences.

In this study, we introduced a method for representing transcriptome components using low-dimensional Raman LDA axes based on Raman-transcriptome linear correspondence (Fig. 4). This method is particularly useful when the Raman LDA axes are closely associated with particular phenotypic characteristics. In our study, the major LDA axes (LDA1 and LDA2) were related to vancomycin susceptibility and virulence (Fig. 1D and 1F). It should be noted that the correspondence between the LDA axes and the phenotypes depends on the presence of systematic differences in their molecular profiles associated with phenotypes. In other words, proximity of different samples along a Raman LDA axis corresponding to some phenotype indicates the presence of shared molecular profile changes related to the phenotype across those samples.

This transcriptome representation facilitates identifying gene groups whose expression patterns correlate with different phenotypes. For instance, we identified several genes expressed in both clinically isolated and laboratory-constructed VISA strains (Table 1 and Fig. 4). Some of these genes have previously been shown to exhibit increased expression levels in VISA strains (*48,49*). However, their role in enhancing vancomycin resistance has not been characterized in detail. Furthermore, some of the identified genes are completely functionally unknown. Characterizing the roles of those genes is an important topic of future studies. Additionally, it would be important to investigate whether only some of these genes contribute to resistance or if the co-expression of these multiple genes is indispensable.

The Mu3 strain was found to be phenotypically heterogeneous in both growth and Raman spectra. This result suggests that cellular Raman spectra can interrogate cell-to-cell heterogeneity within a cellular population. Such phenotypic heterogeneity could be linked to resistance of a subpopulation in the Mu3 strain under vancomycin exposures.

While our study focused on vancomycin resistance in MRSA, the broader applicability of this method to detect drug resistance in other bacterial and fungal pathogens represents an exciting avenue for future research. Exploring the adaptability of Raman spectroscopy to various pathogenic strains could significantly enhance our capacity to combat antibiotic resistance.

## MATERIALS AND METHODS

### Bacterial strains

Nine *S. aureus* strains were used in this study. Three strains, N315, Mu3, and Mu50, are clinical isolates (*5*). The other six strains, ΔIP, ΔIP1, ΔIP2, ΔIP3, ΔIP4, and ΔIP5, were constructed from N315 by introducing mutations or by selection under rifampin exposures (*16*). The vancomycin MIC values of the strains were determined by Etest in (*16*). The vancomycin resistant phenotypes of the strains were determined in (*16*) on the basis of MIC values and population curves on a vancomycin concentration-CFU plane: VSSA for N315 and ΔIP; hVISA for Mu3, ΔIP1, and ΔIP2; and VISA for Mu50, ΔIP3, ΔIP4, and ΔIP5.

### Raman measurement and preprocessing of Raman signals

To prepare samples for Raman measurements, *S. aureus* cells were inoculated from a glycerol stock into 3 mL BHI medium (Eiken Chemical Co., Ltd.) and cultured with shaking (Taitec BR-32FP, 200 r min^−1^) at 37 °C overnight. A 30 µL overnight culture was inoculated into 3 mL BHI medium and grown to an exponential phase (OD_600_ = 2.0–2.5) with shaking at 37 °C. 1 mL of cell suspension was taken in an eppentube, and the cells were washed with phosphate-buffered saline (PBS) twice. The cells were fixed with 4% (w/w) paraformaldehyde for 15 min at room temperature. The fixed cells were washed twice with 0.1M glycine. After substituting the liquid by 1 mL PBS, the cell suspensions were stored at 4 °C until Raman measurement (max. one day).

Immediately before the Raman measurement, the liquid of cell suspension was substituted by 1 mL distilled water. 5 µL of the cell suspension was placed on a synthetic quartz slide glass (TOSHIN RIKO Co., LTD.) and dried.

Raman spectra of the cells were measured using a custom-built Raman microscope (STR-Raman, AIRIX corp.) with 532 nm excitation wavelength (Gem 532, Laser Quantum). Laser power at the sample plane was approximately 17.5 mW. Raman spectra were obtained from single cells with exposure time of 5 s. For each strain, 16 cells were measured per replicate.

Three biological replicates were measured on different days. Consequently, we obtained Raman spectra from 48 cells for each strain. Background Raman spectra were also measured in regions adjacent to target cells. Raman signals were detected by an sCMOS camera (ORCA-Flash4.0 V2, Hamamatsu Photonics) water-cooled at 15 °C. The optical setup was the same as that used for the Raman measurement of *Escherichia coli* in (*33*).

An sCMOS-specific noise reduction filter developed in (*33*) was applied to the obtained images. Then, the region where the spectra were recorded was cropped, and pixel counts were summed along the direction perpendicular to the wavenumber. Next, background Raman spectra were subtracted from single-cell Raman spectra, and the region of 632 to 1862 cm^−1^ was cropped. Subsequently, each spectrum was normalized by subtracting the average and dividing it by the standard deviation.

### Dimension reduction of cellular Raman spectra

Principal component analysis (PCA) was first applied to the normalized spectra to reduce noise. Principal components that explained 98 % of the variance were used. After PCA, linear discriminant analysis (LDA) was applied. LDA is a supervised method, which calculates the most discriminatory bases by maximizing the ratio of the between-group (in the case of this research, between-strain) variances to the sum of within-group (within-strain) variances. The dimension was reduced to *n* − 1 by LDA, where *n* (= 9) was the number of groups (strains).

### Linearly estimating dimension-reduced Raman spectra from transcriptome profiles

Since the number of genes is much larger than that of the strains, and in general, expression patterns of many genes are highly correlated, a linear regression method called partial least squares regression (PLS-R) was employed. The transcriptomes of the nine *S. aureus* strains were obtained from the public database (NCBI GEO, accession number GSE43643, GSE42218, and GSE50826). The dimensions of the transcriptomes were reduced to *n*−3 in PLS-R. To verify the linearity between the LDA Raman spectra and transcriptomes, leave-one-out cross-validation (LOOCV) was performed. We calculated the predicted residual error sum of squares (PRESS) as estimation errors. Statistical significance of the estimation was tested with permutation test. See (*33*) for the detail of LOOCV and permutation test.

### Linearly inferring global transcriptome profiles from dimension-reduced Raman sepctra

To investigate whether global transcriptome profiles can be inferred from the low-dimensional LDA Raman spectra, ordinary linear regression was conducted. We verified the linear correspondence by LOOCV again. In the case of LOOCV, the regression is underdetermined, and the minimum norm solution among all least-squares solutions was adopted. Overall estimation errors were calculated with some distance metrics including the predicted residual error sum of squares (PRESS). Statistical significance of the inference was examined with permutation test.

### Mapping transcriptome profiles on dimension-reduced Raman space

Linear transformation between the transcriptome profiles and the dimension-reduced LDA Raman profiles is

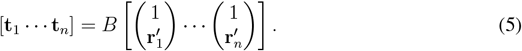

As 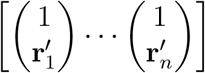 is full rank, the strain-independent coefficient matrix *B* can be obtained by

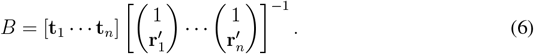

Writing the *j*-th column of *B* as **b**_*j*_, *B* = [**b**_0_ *·· ·* **b**_*n*−1_]. From the viewpoint of linear regression, **b**_0_ is the constant terms, and **b**_*j*_ (1 ≤ *j* ≤ *n* − 1) is the coefficients for the *j*-th LDA dimension. The normalized coefficients discussed in the main text is the columns of diag (**b**_0_)^−1^ *B*, where diag (**x**) = (*δ*_*ij*_*x*_*i*_) for a vector **x** = (*x*_*i*_). For instance, the first and second axes in Fig. 4 are the second and third axes of diag (**b**_0_)^−1^ *B*, namely, diag (**b**_0_)^−1^ **b**_1_ and diag (**b**_0_)^−1^ **b**_2_. See (*45*) for the detailed mathematical explanation of the phenotype-expression pattern correspondence between the LDA Raman space and the normalized coefficient gene space.

### Heterogeneity analysis of Raman spectra of the clinically isolated strains

To make the intensities of the average spectra approximately equal among the three strains, three outlier single-cell Raman spectra were removed for each strain. Then, the standard deviation of the remaining 45 normalized spectra was calculated at each wavenumber for each strain.

### Time-lapse microscopy

To prepare samples for time-lapse microscopy, *S. aureus* cells were inoculated from a glycerol stock into 2 mL BHI medium and cultured with shaking (200 r min^−1^) at 37 °C overnight. A 15 µL overnight culture was inoculated into 3 mL BHI medium and grown to exponential phase (OD_600_ = 2.4–2.5) with shaking at 37 °C. 1 µL of cell suspension was taken on a coverslip (Neo No.1, 24 mm*×*60 mm, Matsunami) and covered by a BHI agarose pad (20 mm*×*25 mm*×*7 mm (thickness)) solidified with 1.5% low-melt agarose (GeneMate LowMelt Agarose, BioExpress). The assembled sample was placed on a microscope XY stage and covered by a transparent plastic dish to prevent the agarose from drying.

We used Nikon Eclipse Ti-E inverted microscope equipped with digital sCMOS camera (ORCA-Flash4.0 V3, Hamamatsu Photonics), phase-contrast objective (Plan Apo *λ*, 100*×*, NA1.45). We obtained phase-contrast images from 50 XY positions with 1 min interval in each experiment.

The acquired time-lapse images were analyzed using ImageJ (*57*). We applied Mexican Hat Filter to find outlines of microclonies of *S. aureus* and measured the areas to quantify the growth.

### Heterogeneity analysis of microcolony growth of the clinically isolated strains

Growth rates were calculated by taking logarithm to 2 of relative colony sizes at *t* = 3 h to those at *t* = 0h and dividing them by 3 h.

## Data analysis

All the data analyses in this study were conducted using MATLAB (R2019a, R2023b and R2024b).

## Acknowledgments

We thank Tetsuo Yamaguchi for reading the manuscript and providing comments. This work was supported by JST CREST Grant Number JPMJCR1927 (Y.W.); JST ERATO Grant Number JPMJER1902 (Y.W.); JSPS KAKENHI Grant Numbers 19J22448 (K.F.K.); and Takeda Science Foundation Award (Y.K. and Y.W.).

---

**Table 1.** List of genes with a high *b*_*i*1_*/b*_*i*0_ value. Genes with a *b*_*i*1_*/b*_*i*0_ value above 1.5, which are located on the right side and highlighted by red circles in Fig. 4, are listed.

