## Supplemental text and figures for "Raman spectra identify vancomycin-resistant phenotypes and their transcriptomic features in *Staphylococcus aureus*"

### Supplementary Materials

**Figure S1 Single-cell Raman spectra. Related to Fig. 1. (A)** Schematic figure of the Raman microscope used in this study. **(B)** Representative single-cell Raman spectra of the remaining seven strains not shown in Fig. 1C.

**Figure S2 Strong correlation between transcriptome profiles of different strains. Related to Fig. 3.** Scatterplots between the measured transcriptome of N315 and the measured transcriptomes of the other strains. The transcriptome data were obtained from public databases (*1–3*) (Materials and Methods).

**Table S1 The overall prediction errors of transcriptomes with various distance metrics. Related to Fig. 3.** The overall estimation errors of transcriptomes from Raman spectra were evaluated with various distance measures. $p$-values of permutation test (1,000 permutations) were shown. With all the metrics, $p$-values were $9.99 \times 10^{-4}$, which mean that no permuted dataset produced a prediction error lower than that of the original dataset.

**Movie S1 Time-lapse movie of microcolony growth of N315.** Time stamp shows hour:min.

**Movie S2 Time-lapse movie of microcolony growth of Mu3.** Time stamp shows hour:min.

**Movie S3 Time-lapse movie of microcolony growth of Mu50.** Time stamp shows hour:min.

### Fig. S1

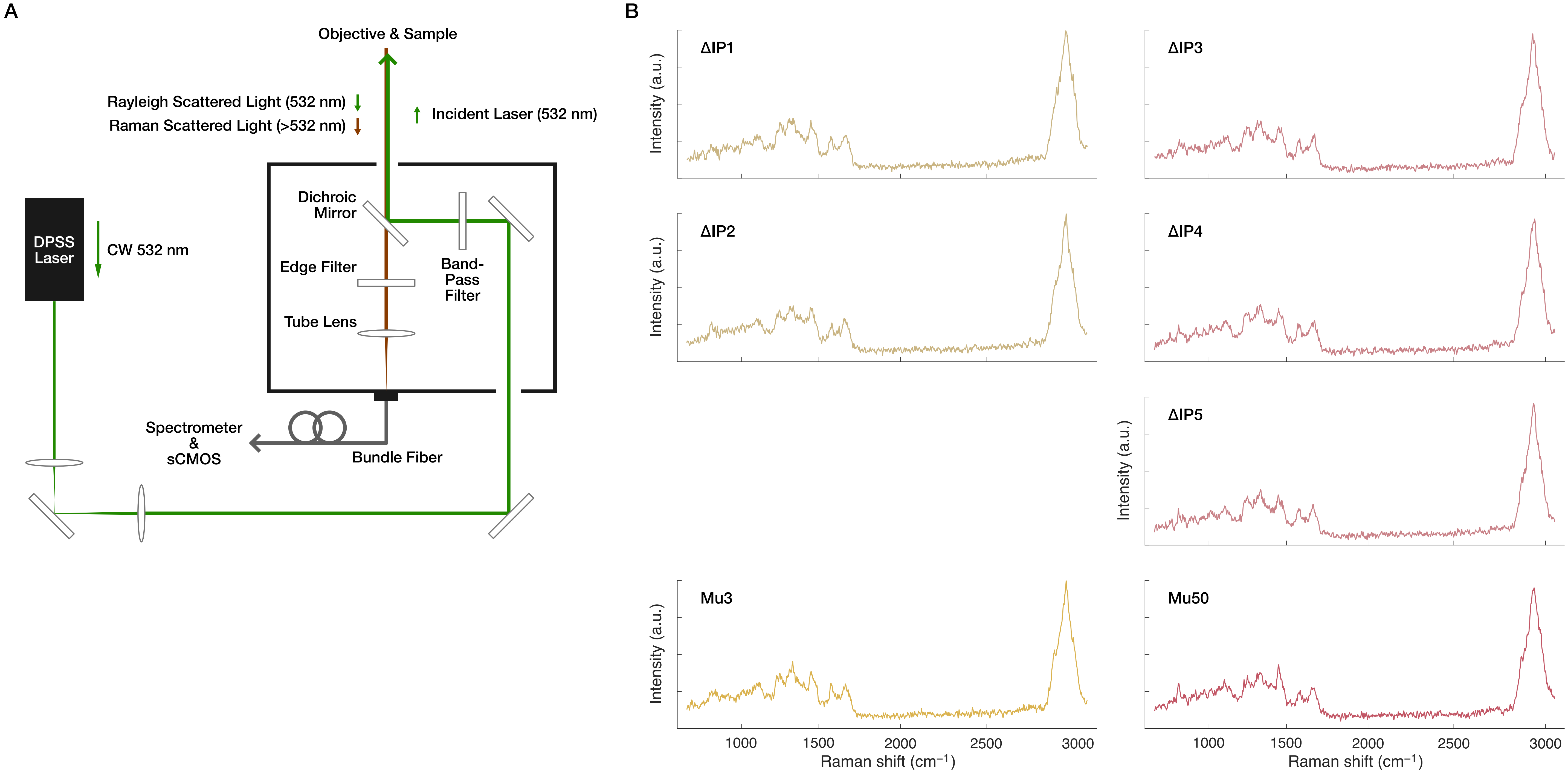
A
Objective & Sample
Rayleigh Scattered Light (532 nm)
Raman Scattered Light (>532 nm)
Incident Laser (532 nm)
DPSS Laser
CW 532 nm
Dichroic Mirror
Edge Filter
Band-Pass Filter
Tube Lens
Spectrometer & sCMOS
Bundle Fiber
B
ΔIP1
ΔIP2
ΔIP3
ΔIP4
ΔIP5
Mu3
Mu50
Intensity (a.u.)
1000
1500
2000
2500
3000
Raman shift (cm−1)

### Fig. S2

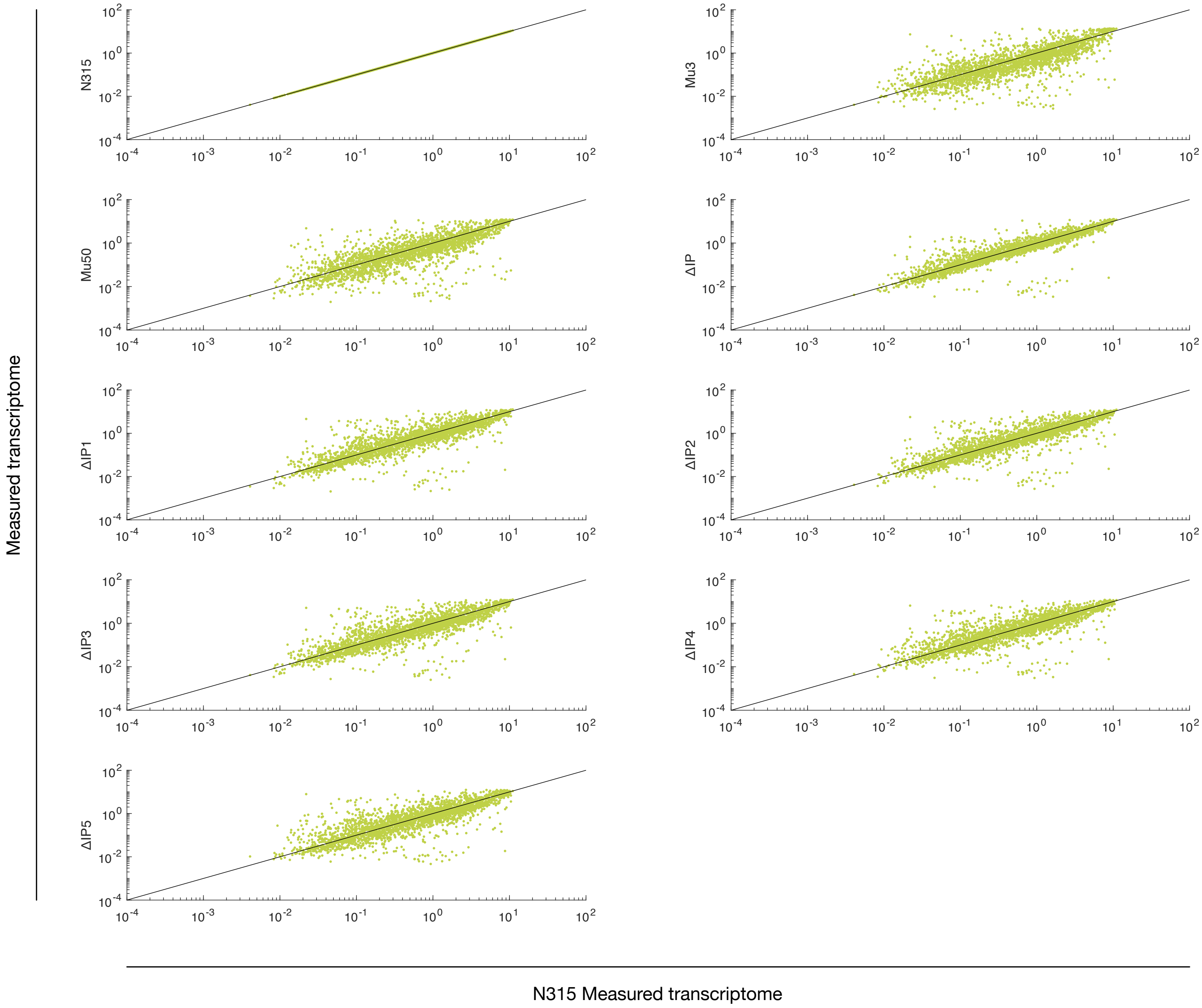
N315
Mu3
Mu50
ΔIP
ΔIP1
ΔIP2
ΔIP3
ΔIP4
ΔIP5
Measured transcriptome
N315 Measured transcriptome
